# A centronuclear myopathy-causing mutation in dynamin-2 perturbs the actin-dependent structure of dendritic spines leading to excitatory synaptic defects in a murine model of the disease

**DOI:** 10.1101/2021.06.28.450172

**Authors:** Jorge Arriagada-Diaz, Bárbara Gómez, Lorena Prado-Vega, Michelle Mattar-Araos, Marjorie Labraña-Allende, Fernando Hinostroza, Ivana Gajardo, María José Guerra-Fernández, Jorge A. Bevilacqua, Ana M. Cárdenas, Marc Bitoun, Alvaro O. Ardiles, Arlek M. Gonzalez-Jamett

## Abstract

Dynamin-2 is a large GTP-ase, member of the dynamin superfamily, that regulates membrane remodeling and cytoskeleton dynamics. In the mammalian nervous system dynamin-2 modulates synaptic vesicle (SV)-recycling at the nerve terminals and receptor-trafficking to and from postsynaptic densities (PSDs). Mutations in dynamin-2 cause autosomal dominant centronuclear myopathy (CNM), a congenital neuromuscular disorder characterized by progressive weakness and atrophy of distal skeletal muscles. Cognitive defects have also been reported in dynamin-2-linked CNM patients suggesting a concomitant impairment of the central nervous system. Here we addressed the mechanisms that lead to cognitive defects in dynamin-2-linked CNM using a knock-in mouse model that harbors the p.R465W mutation in dynamin-2, the most common causing CNM. Our results show that these mice exhibit reduced capability to learn and acquire spatial and recognition memory, impaired long-term potentiation of the excitatory synaptic strength and perturbed dendritic spine morphology, which seem to be associated with actin defects. Together, these data reveal for the first time that structural and functional synaptic defects underlie cognitive defects in the CNM context. In addition our results contribute to the still scarce knowledge about the importance of dynamin-2 at central synapses.

## Introduction

Dynamin super-family (DSF) is a group of large GTP-ases that act as mechano-enzymes promoting membrane remodeling in several processes including exocytosis, endocytosis, intracellular trafficking and mitochondrial dynamics among others (Ferguson, et al., 2012; González-Jamett, et al., 2013; Lomash, et al., 2015; Arriagada-Diaz, et al., 2020). Classical dynamins are the best understood members of the DSF. In mammals, these are encoded by three different genes: *DNM1, DNM2* and *DNM3* located on chromosomes 9, 19 and 1, respectively (Newman-Smith, et al., 1997; Züchner, et al., 2005; Noakes, et al., 1999). Classical dynamins are composed by five highly conserved domains: an amino-terminal GTP-ase domain that binds and hydrolyze GTP, a middle structural domain, a PH-domain involved in lipid membrane interaction, a GTP-ase effector domain (GED) and an arginine- and proline-rich domain (PRD) that allows dynamin association with SH3-containing proteins (Praefcke, et al., 2004; Ferguson, et al., 2012; Antonny, et al., 2016; Singh, et al., 2017; Arriagada-Diaz, et al., 2020). These domains organize in three regions: a “bundle signaling element” (BSE) composed by helices from the GTP-ase domain and GED, a “stalk” composed by helices from the middle domain and GED, and a membrane-inserting “foot” formed by the PH domain (Chappie, et al., 2009; Faelber, et al., 2011; Ford, et al., 2011; Kong, et al., 2018). In response to the binding of GTP, the BSE convey conformational changes from the GTP-ase domain to the “stalk” promoting dynamin oligomerization in helical structures that enhance its GTP-ase activity and favor membrane scission (Hinshaw, et al., 1995; Warnock, et al., 1997; Antonny, et al., 2016). Dynamin assembly/disassembly cycles and GTP-ase activity also regulate the bundling of the actin cytoskeleton (Gu, et al., 2010; Zhang, et al., 2020; Lin, et al., 2020; Schiffer, et al., 2015a).

The three classical dynamins are expressed at the central nervous system (CNS) and play functions at the pre- and post-synapses (Arriagada-Diaz, et al., 2020). At the pre-synapses they support the endocytic recycling of synaptic vesicles (SVs), exerting differential roles depending on the pattern of neuronal activity (Tanifuji, et al., 2013). In this regard, dynamin-1 mediates SV recycling upon high-frequency stimulation but responding in a slow time window after the arrival of action potentials (APs); dynamin-3 works in a faster time window independent of the frequency of APs and dynamin-2 mediates SV resupply upon high-frequency stimulation with a fast kinetics (Tanifuji, et al., 2013).

At the postsynaptic level dynamin-2 is an important regulator of surface-membrane availability of neurotransmitter-receptors (Carroll, et al., 1999; Kabbani, et al., 2004; Bhatnagar, et al., 2000; Wang, et al., 2017). The latter appears to be especially relevant for glutamatergic excitatory synapses, as dynamin-2 modulate insertion (Jaskolski, et al., 2009), removal (Carroll, et al., 1999; Chowdhury, et al., 2006), and recycling (Lu, et al., 2007; Zheng, et al., 2015) of the α-amino-3-hydroxy-5-methyl-4-isoxazolepropionic acid (AMPA) receptor to and from PSDs.

Although the isoforms -1 and -3 are expressed at a more greater level than dynamin-2 at the CNS (Okamoto, et al., 2001), dynamin-2 seems to play a more critical role for synapse development (Ferguson, et al., 2009). In fact, while embryos of dynamin-1 and/or dynamin-3 knockout (KO) mice survive for weeks after born (Ferguson, et al., 2007; Raimondi, et al., 2011), the KO of dynamin-2 results in early embryonic lethality (Ferguson, et al., 2009).

Mutations in dynamin-2 cause autosomal dominant human diseases (Züchner, et al., 2005; Bitoun, et al., 2005; Tanabe, et al., 2009; Liu, et al., 2011; Koutsopoulos, et al., 2011; Böhm, et al., 2012; Sambuughin, et al., 2015; Ali, et al., 2019). Among them, mutations in the middle and PH domain of dynamin-2 cause the autosomal dominant form of centronuclear myopathy (CNM) a rare neuromuscular disorder clinically manifested by myalgia, fatigability, weakness, and progressive atrophy of distal skeletal muscles (Jeannet, et al., 2004; Fischer, et al., 2006; Böhm, et al., 2012). Cognitive deficiencies, manifested as learning disabilities and limited intelligent quotient, have also been reported in CNM patients (Jeannet, et al., 2004; Fischer, et al., 2006; Echaniz-Laguna, et al., 2007; Böhm, et al., 2012) although the mechanisms involved have never been described. Here we show that a knock-in (KI) mouse model bearing the CNM-causing p.R465W mutation in dynamin-2 (Durieux, et al., 2010; Durieux, et al., 2012) exhibit a deficient capability to learn and acquire spatial and recognition memory. These cognitive defects correlate with impaired hippocampal excitatory synaptic transmission and plasticity. Furthermore, reduced dendritic spine density and changes in spine morphology are also evident in the brain of these mice. This latter seem to rely on defects in the actin cytoskeleton organization. As the catalytic activity of dynamin-2 is an important regulator of the actin dynamics (Mooren, et al., 2009; Gu, et al., 2010; Yamada, et al., 2013; Yamada, et al., 2016; Lin, et al., 2020) and this is a mechanism affected in dynamin-2 linked CNM (González-Jamett, et al., 2017) these data strongly suggest that CNM-causing mutations perturb the synaptic role of dynamin-2, impacting on the actin-dependent dendritic-spine structure and consequently affecting synaptic transmission and cognitive functions.

## Results

### Impaired learning and memory in adult CNM-mice

As early learning disabilities and cognitive defects have been reported in dynamin-2-related CNM patients (Jeannet, et al., 2004; Fischer, et al., 2006; Echaniz-Laguna, et al., 2007; Böhm, et al., 2012) we first evaluated learning and memory in a murine model of CNM. This is a knock-in mouse harboring the p.R465W mutation located in the middle domain of dynamin-2, the most common mutation causing CNM (Durieux, et al., 2010; Durieux, et al., 2012). Heterozygous mice (HTZ) recapitulate most of the signs of the myopathy, starting at 1 month old (m.o) and progressing as the animal ages (Durieux, et al., 2010). In order to assay learning and memory capabilities, adult HTZ mice over 6 m.o and their age-matched wild type (WT) littermates were subjected to two different behavioral paradigms: the novel object recognition test (NOR) and the Barnes maze (BM). Whilst NOR is useful to evaluate recognition memory (Ennaceur, et al., 1988), BM is used to measure the ability of mice to learn and remember a spatial location (Rizzo, et al., 2017; Negrón-Oyarzo, et al., 2015; Pitts, et al., 2018).

First, we performed the NOR test. A schematic representation of the test is shown in Figure 1a. As expected WT mice spent significantly more time exploring a novel object (N) than a familial object (F) along sessions (Figure 1b). However HTZ mice did not modify their preference between N and F (Figure 1b) suggesting defects in the recognition memory. Moreover, HTZ mice exhibited a significantly reduced recognition index (D1) compared to their WT littermates (Figure 1c) strongly suggesting a decline in the recognition memory in the CNM context.

**Figure 1:**
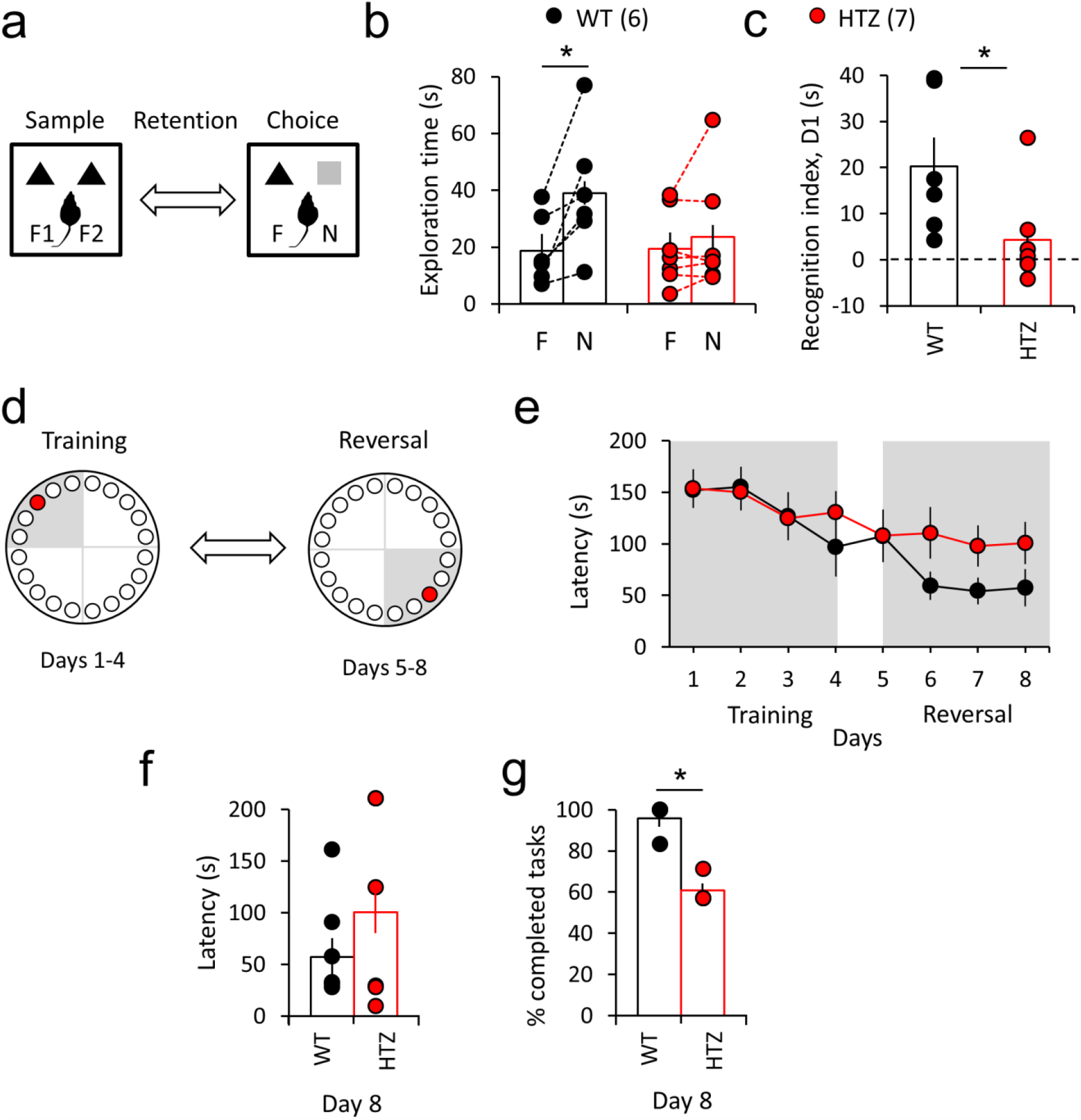
HTZ mice exhibit reduced performance in recognition and spatial memory tasks. WT (black circles) and HTZ (red circles) mice over 6 m.o were evaluated in the novel object recognition (NOR) and the Barnes maze tests to estimate recognition and spatial memory, respectively. **(a)** Representative scheme of the NOR test. Mice are faced with two identical objects (F1 and F2) in the sample phase and then in the choice phase, the objects are replaced by an extra copy of the familiar object (F) and a novel object (N). **(b)** Mean exploration time for familial (F) and novel (N) objects. Note that HTZ mice spend significantly less time exploring new objects compared to WT mice. Wilcoxon test, *p =0.0312 compared to the WT condition. **(c)** Mean recognition index (D1). Positive values indicate recognition of the novel over the familial object. Mann-Whitney test, *p =0.0221 compared to the WT condition **(d)** Schematic representation of the BM test. Mice were trained for 4 days to find and escape to a hidden dark box using visual cues (Training phase). On the fifth day, the escape box is located in the opposing quadrant keeping the visual cues to evaluate spatial memory flexibility (Reversal phase). **(e)** Quantification of the latency to entry into the escape box throughout the training and reversal sessions. **(f)** Mean latency to entry into the escape box at day 8 of the test (last day of the reversal phase). **(6)** Quantification of the percentage of animals that completed the task at the end of the reversal phase. Note that the the time to enter into the escape box was not statistically significant, but the percentage of HTZ mice that completed the task on the last day of the reversal phase was significantly lower than WT. Mann-Whitney test, *p =0.0286 compared to the WT condition All data are represented as mean ± SEM. In parentheses is the number of WT and HTZ mice.

Then we applied the BM test. A schematic representation of the test is shown in Figure 1d. WT and HTZ mice were trained during 4 days (training) to find and enter into a hidden scape box using spatial clues. On the fifth day a reversal phase started in which the position of the scape-box was changed but the visual clues were kept to evaluate memory flexibility. As shown in Figure 1e the latency to find and enter into the scape box was decreasing throughout the training sessions for both WT and HTZ mice. However, HTZ mice took a longer time than WT to access the scape box throughout the reversal phase (Figure 1f). In fact, on the last day of the reversal phase WT mice took on average 57.3 ± 18.2 s whereas, HTZ mice took 100.6 ± 20.5 s to end the test (Figure 1f). Remarkably the percentage of HTZ mice that completed the test at the last day of the reversal phase was significantly lower than WT mice (Figure 1g) suggesting defects in spatial memory flexibility. Importantly, these observations did not appear to be due to locomotion defects in HTZ mice as walking time percentage (Figure supplement S1a) and total distance traveled in an open-field arena (Figure supplement S1b) were not different between groups.

Together these data show that the DNM2-linked CNM mouse model shares cognitive impairment with CNM patients and represents a pertinent model to identify the underlying molecular mechanisms

### Excitatory synaptic transmission and plasticity are defective in adult hippocampal slices from CNM-mice

Changes in neuronal activity induce modifications in the efficacy of the synaptic transmission; this is known as synaptic plasticity and the mechanisms regulating it are the basis of learning and memory (Shors, et al., 1997). As synaptic plasticity in the hippocampus is particularly important for memory formation (Lee, et al., 2011) and the most studied forms of synaptic plasticity are long-term potentiation (LTP) and long-term depression (LTD) of the synaptic strength at the CA3-to-CA1 synapses (Malenka, et al., 2004) we evaluated excitatory synaptic transmission and plasticity in hippocampal slices isolated from HTZ mice over 6 m.o and their age-matched WT littermates.

The strength of the basal excitatory synaptic transmission was estimated as the I/O relationship. Figure 2a schematizes the experimental configuration for electrophysiological field recordings in hippocampal slices. Figure 2b show representative I/O traces for WT and HTZ hippocampal slices. As shown in Figure 2c the I/O relationship was significantly impaired in HTZ compared to WT slices. HTZ slices exhibited significantly lower fEPSP slopes (Figure 2d) but significantly higher FV-amplitudes (Figure 2e) compared to the WT condition. To evaluate whether plasticity of the synaptic strength is also affected in the CNM-context we induced NMDAR-dependent LTP in WT and HTZ hippocampal slices by using a standard theta burst stimulation (TBS) protocol that drives a compound-LTP, involving pre- and post-synaptic mechanisms (Bayazitov, et al., 2007). Whilst LTP induction relies on the activation of NMDAR-that requires both presynaptic glutamate release and post-synaptic depolarization (Citri, et al., 2008), LTP maintenance mainly relies on AMPAR trafficking at the PSDs (Anggono, et al., 2012a). As shown in Figure 3 the magnitude of the TBS-induced LTP was significantly reduced after 50 min of record in HTZ compared to WT hippocampal slices (Figures 3a and 3c) suggesting a role of dynamin-2 in the maintenance of the tetanus-induced LTP. Furthermore, the early potentiation of the synaptic response measured during the first 5 minutes after the application of TBS was also reduced in HTZ compared to WT slices (Figure 3b) suggesting that LTP-induction is also dependent on dynamin-2. To evaluate the presynaptic contribution to the defects observed in HTZ synapses we estimated a paired-pulse facilitation (PPF) index (Figure 3d-e). PPF is a form of short-term synaptic plasticity that occurs when two stimulation-pulses of similar intensities are given in rapid succession, causing the second synaptic response to be bigger than the first one (Zucker, et al., 2002). Facilitation appears to be due to an increased presynaptic Ca^2+^ concentration that leads to enhanced neurotransmitter-release probability (Zucker, et al., 2002). The PPF index was unchanged between HTZ and WT hippocampal slices (Figure 3d-e) suggesting that, although presynaptic mechanisms could operate in the CNM-context, those appear to be independent of the release probability. Hence, the p.R465W mutation in dynamin-2 seems to mainly affect post-synaptic mechanisms, impairing synaptic potentiation upon sustained neuronal activity. The main post-synaptic impact was confirmed by a chemically induced LTP using the NMDAR-agonist Glycine (chLTP) which activates NMDAR without mediating a presynaptic glutamate release (Lu, et al., 2001; Zhang, et al., 2014). Upon chLTP a significant reduction in the magnitude of synaptic potentiation was observed in HTZ compared to WT slices (Figure supplement S2). Interestingly, these postsynaptic mechanisms appear to predominantly affect excitatory synaptic transmission, as we observed similar differences between HTZ and WT slices when the TBS protocol was elicited in the presence of *picrotoxin* (*PTX*), a blocker of the GABAR-dependent inhibitory transmission (Figure supplement S3).

**Figure 2:**
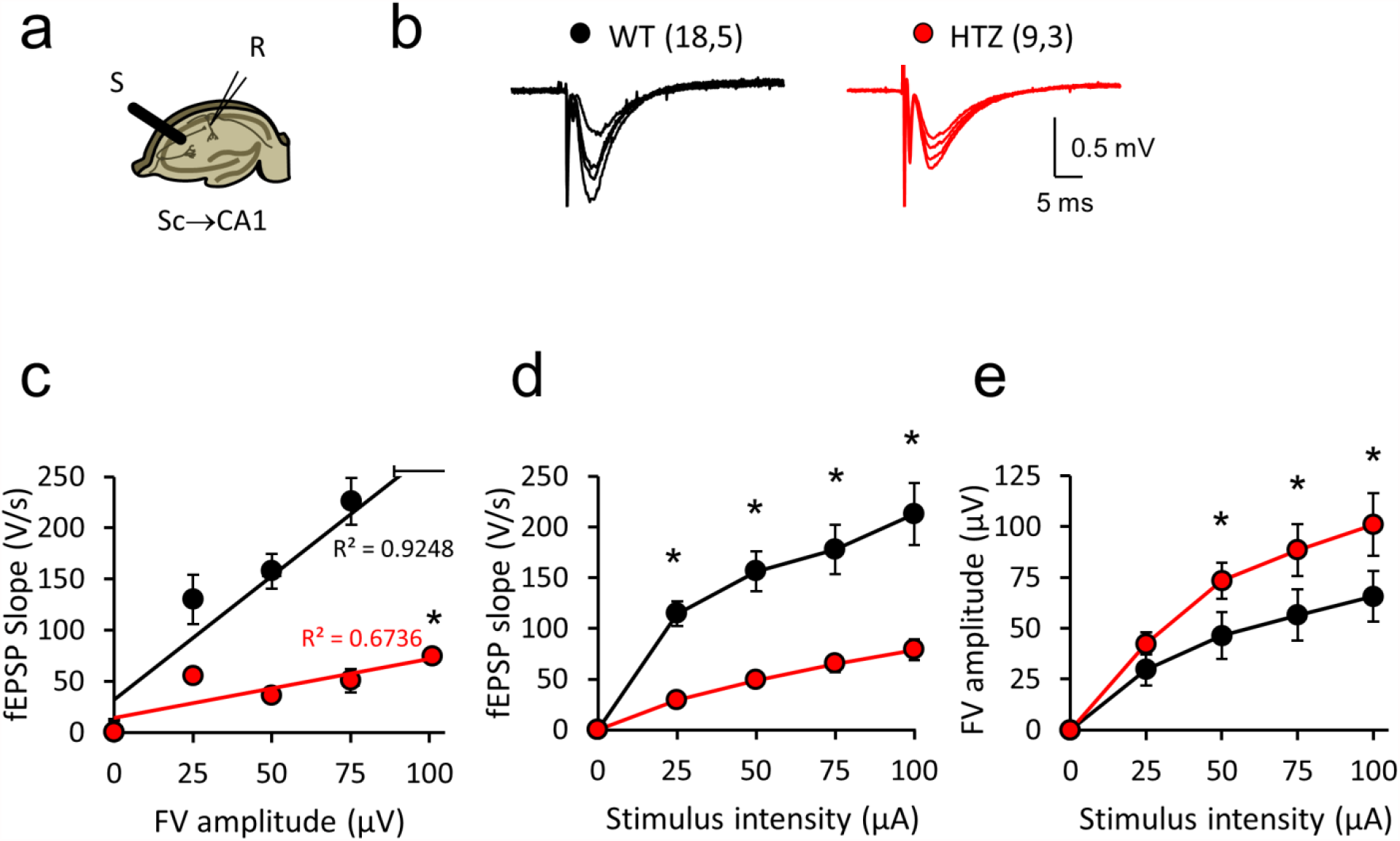
Impaired basal excitatory synaptic transmission in HTZ hippocampal slices **(a)** Scheme of the experimental configuration for field recordings in Schaffer Collaterals (Sc) to CA1 hippocampal synapses. The stimulation electrode and the recording electrode (R) were positioned in *stratum radiatum* of CA1. **(b)** Representative fEPSP traces of both experimental groups HTZ (red) and WT (black). The numbers in parentheses are slices, animals recorded. **c)** Relationship between the slope values of fEPSPs and the amplitude of FVs obtained between each experimental groups. The R^2^ values for the respective linear regressions are shown. **(d)** Mean fEPSP slopes at different intensity values are plotted per each experimental group. Note the significant difference in fEPSPs as the intensity of stimulation increased between WT and HTZ slices. **(e)** Mean FV amplitudes for WT and HTZ slices. Note the significant difference in FV amplitudes between WT and HTZ slices as the stimulus intensity increased. (WT: n = 18 slices, 5 mice; HTZ: n = 9 slices, 3 mice). All data are represented as mean ± SEM. Statistical differences were calculated using a t-test, * p <0.0001 compared to the WT condition.

**Figure 3:**
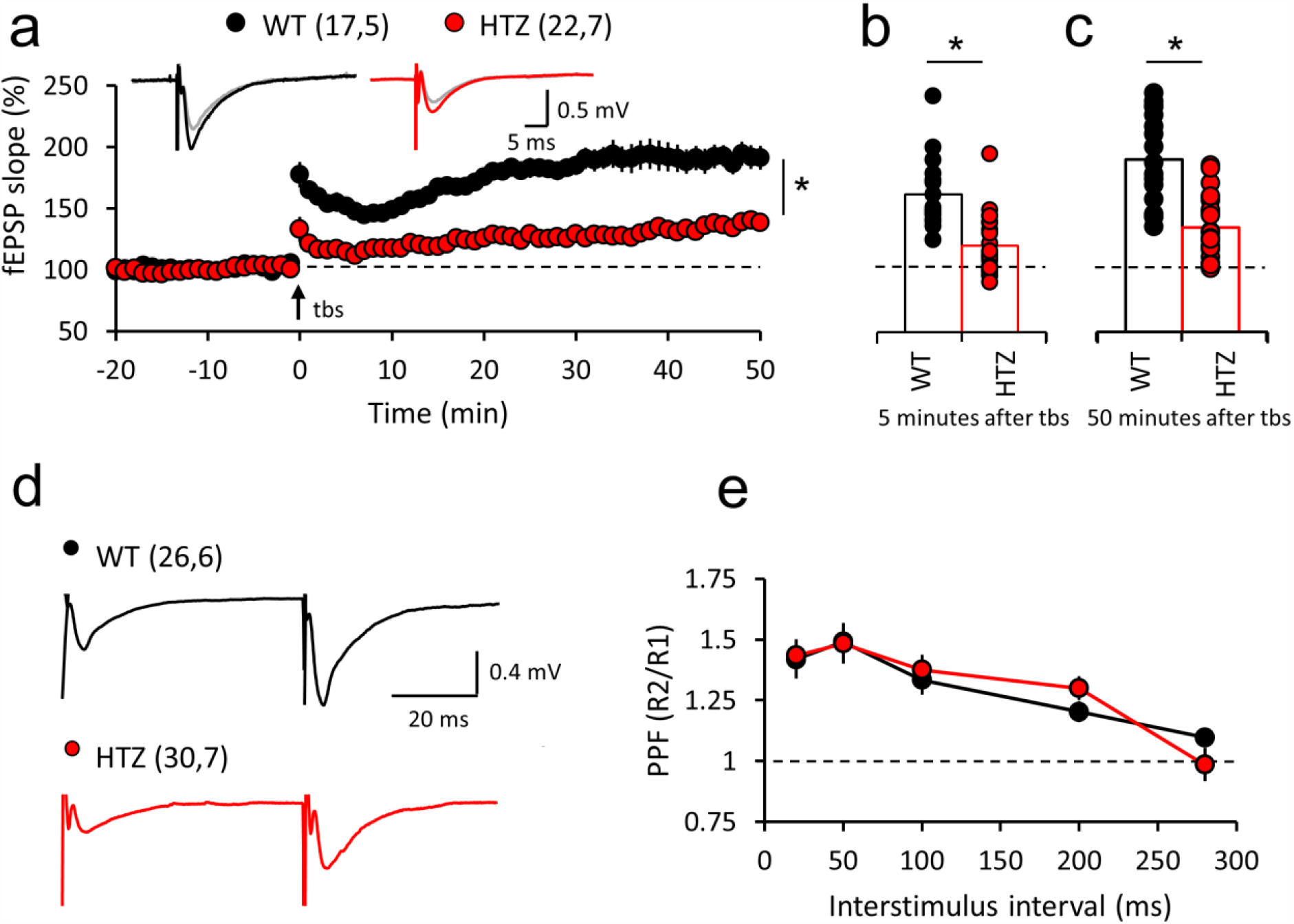
TBS-induced LTP impairment in excitatory synapses of HTZ hippocampal slices. **(a-c)** Hippocampal slices of 6 m.o of WT (black circles) and HTZ (red circles) mice were stimulated with a standard TBS protocol to induce LTP and excitatory postsynaptic field potentials (fEPSP) were recorded during 50 min after the application of the TBS protocol. 2-Way ANOVA, F_(1,2257)_ = 1414, *p <0.0001 followed by Bonferroni post hoc test (p < 0.05) compared to the WT condition. Representative traces before (gray) and after (black and red) the application of TBS are shown as inserts. Note that at the end of the recording (c), as well as after 5 min of LTP-induction (b) HTZ slices exhibited a significantly reduced potentiation of the synaptic strength compared to WT slices (WT: n = 17 slices, 5 mice; HTZ: n = 22 slices, 7 mice). Mann-Whitney test, *p <0.0001 compared to the WT condition in both 5 or 50 min after TBS. The numbers in parentheses are slices, animals recorded **(d)** Representative traces of HTZ (red) and WT (black) slices stimulated with two stimuli of the same intensity separated by a time of 50 ms for PPF index estimation. Note that the amplitude of the second response was greater than the first one in both cases. **(e)** Paired pulse facilitation by different stimulation intervals. Note that no significant differences were observed between the experimental groups (WT: n = 26, slices 6 mice; HTZ: n = 30 slices, 7 mice). All data are represented as mean ± SEM.

Like LTP, LTD is a relevant long-term synaptic plasticity phenomena also implicated in learning and memory (Malenka, et al., 2004; Citri, et al., 2008). We evaluated weather LTD is also affected in the CNM-context and observed a significant reduction in the LTD-magnitude in HTZ compared to WT slices (Figure supplement S4).

Altogether these data demonstrate that the excitatory synaptic transmission and plasticity are DNM2-dependent processes impaired in the CNM context.

### Dendritic-spine morphology is perturbed in neurons from adult CNM-mice

Besides a functional excitatory synaptic plasticity that mainly relies on changes in AMPAR activity at the PSDs (Anggono, et al., 2012b) there is a structural plasticity involving morphological changes at dendritic spines. Dendritic spines are actin-enriched protrusions where occur most of the excitatory synapses (Bosch, et al., 2012). Their density, size and shape correlate with the degree of synapse maturity and functionality and undergo modifications upon synaptic plasticity. In this regard it has been demonstrated that LTP-induction increases dendritic spine density and promotes the transit towards a mature-spine shape with higher head diameter leading to a higher head/spine length ratio (Matsuzaki, et al., 2001; Okamoto, et al., 2009). On the contrary, LTD-induction reduces dendritic spine density and leads to the shrinkage of pre-existing spines (Okamoto, et al., 2004). In order to evaluate whether the defects in excitatory synaptic transmission and plasticity observed in HTZ slices were related to structural defects in dendritic spines, we marked brains from 6 m.o WT and HTZ mice with a Golgi- - staining and quantified dendritic spine density and morphology in hippocampal and cortical neurons. Figures 4 and 5 summarize observations made on hippocampal and cortical pyramidal neurons, respectively. The comparison of the mean morphological parameters revealed a significant reduction in spine density (number of spines per 1 µm of dendritic shaft) in HTZ compared to WT hippocampal neurons (Figure 4b). Although no significant differences were observed in the spine shape on average (Figure 4c) HTZ hippocampal neurons tended to exhibit a lower percentage of filopodial-protrusions and a higher percentage of stubby-like spines. To further evaluate dendritic spine structure, we quantified the cumulative frequency of spine lengths, head diameters and head/length ratios. Remarkably, although head diameters were not different between WT and HTZ hippocampal neurons (Figure 4d) these latter exhibited a significantly higher frequency of shorter spines, producing a leftward shift in the cumulative distribution curve compared to WT spines (Figure 4e). As a consequence of having shorter dendritic spines, HTZ hippocampal neurons also exhibited a significantly higher head diameter/spine length ratio compared to WT neurons (Figure 4f).

**Figure 4:**
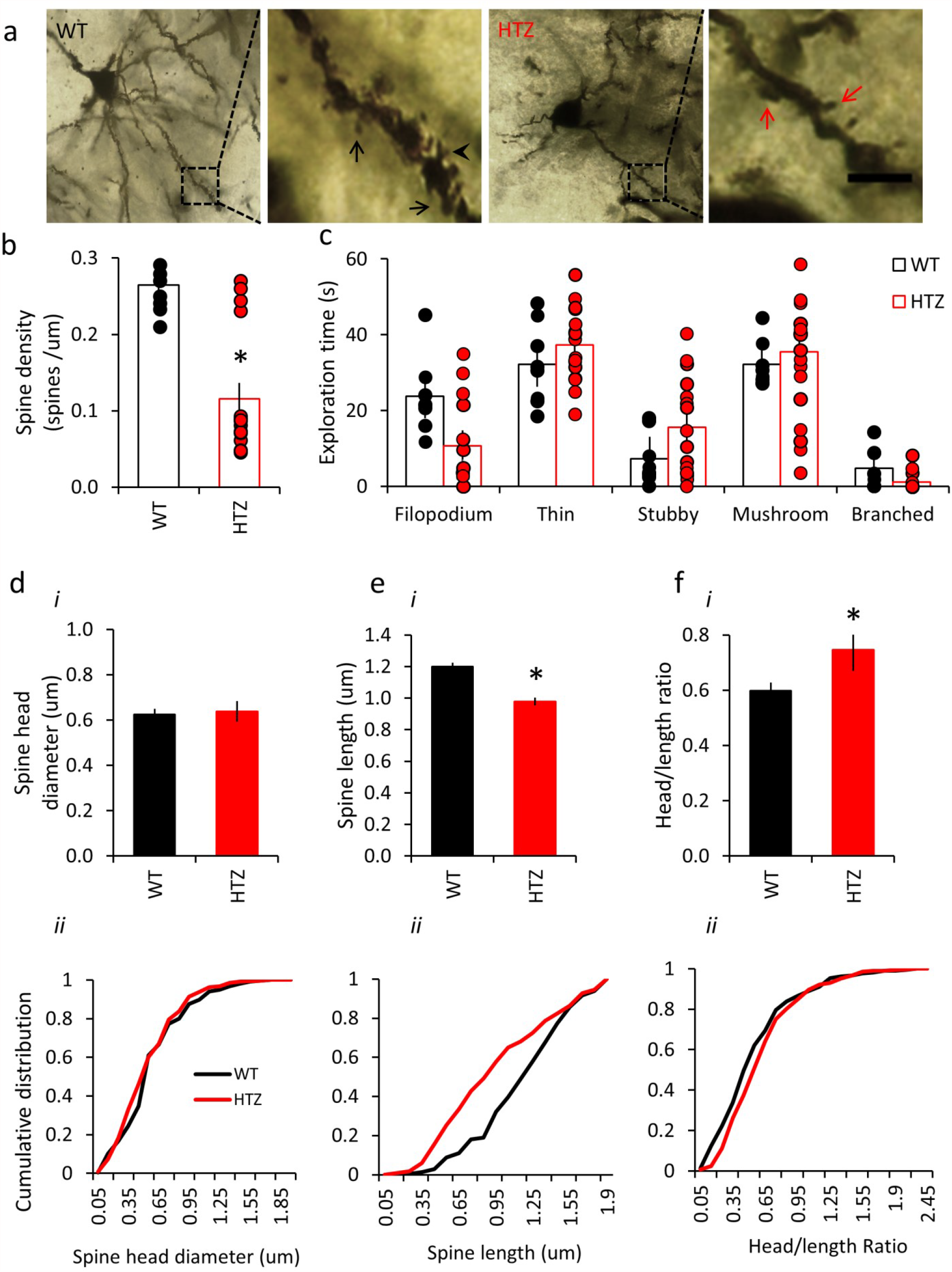
Reduced spine density and spine length in hippocampal neurons from adult HTZ brains. Brains from WT (black bars) and HTZ (red bars) mice over 6 m.o were isolated and Golgi-stained for visualization of pyramidal hippocampal neurons **(a)** Representative neurons and dendritic shafts per genotype. Black and red arrows point spines in WT and HTZ dendrites respectively. Black arrowhead highlights a branched spine. Scale bar= 2.5 µm **(b)** Mean spine density per neuron (number of spines per 1 µm of dendritic segment) **(c)** Mean percentage of filopodium, thin, stubby, mushroom and branched spines in WT and HTZ hippocampal neurons. 4 dendritic shafts per neuron were analyzed. N=8 WT neurons and 19 HTZ neurons from at least 2 different animals per genotype. Bars are mean ± SEM **(d-f)** Average (i) and cumulative distribution (ii) of spine head diameters (d), spine lengths (e) and head/length ratios (f) in WT and HTZ Golgi-stained neurons. N=217 WT spines, 332 HTZ spines. *p<0.005 versus WT condition, Mann-Whitney 2-tail test for non-parametric data.

**Figure 5:**
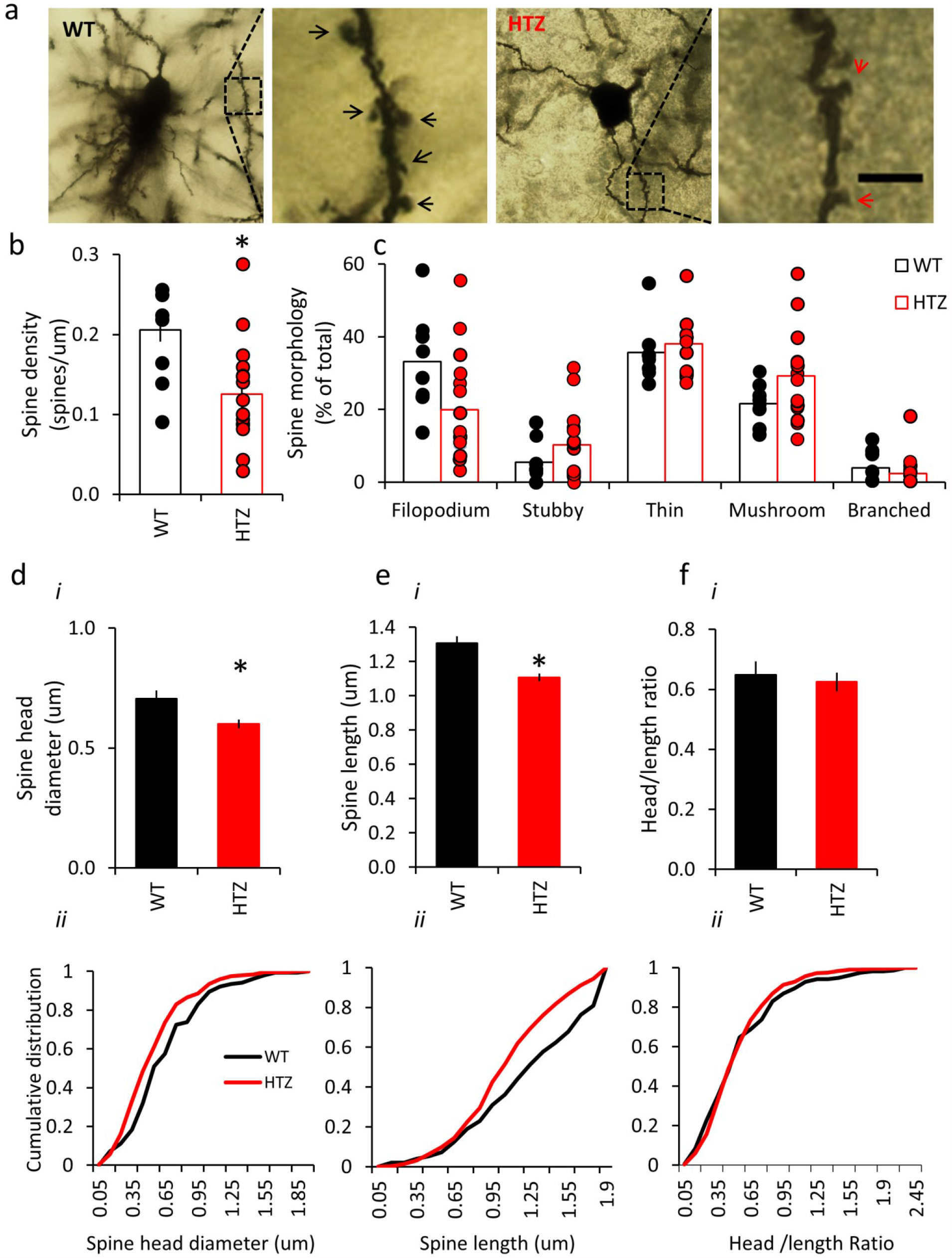
Reduced spine density and spine head diameter in cortical neurons from adult HTZ brains. Brains from WT (black bars) and HTZ (red bars) mice over 6 m.o were isolated and Golgi-stained for visualization of pyramidal cortical neurons **(a)** Representative neurons and dendritic shafts per genotype. Black and red arrows point spines in WT and HTZ dendrites respectively. Scale bar= 2.5 µm **(b)** Mean spine density per neuron **(c)** Mean percentage of filopodium, thin, stubby, mushroom or branched spines in WT and HTZ cortical neurons. 4 dendritic shafts per neuron were analyzed N= 8 WT neurons and 19 HTZ neurons from at least two different animals per genotype. Bars are mean ± SEM **(d-f)** Average (i) and cumulative distribution (ii) of spine head diameters (d), spine lengths (e) and head/length ratios (f) in WT and HTZ Golgi-stained neurons. N=155 WT spines, 328 HTZ spines. *p<0.005 versus WT condition, Mann-Whitney 2-tail test for non-parametric data.

Similar to that observed in hippocampal neurons, HTZ cortical neurons exhibited a significantly lower mean spine density (Figure 5b) and tended to show a lower proportion of filopodial-protrusions compared to the WT condition (Figure 5c). Remarkably, cortical HTZ neurons exhibited a higher frequency of spines with smaller heads (Figure 5d) and shorter lengths (Figure 5e) although the head/length ratio was unchanged (Figure 5f).

Together, these data suggest that pyramidal neurons bearing the dynamin-2 p.R465W mutation may undergo modifications in density, head width and length of dendritic spines.

### Impaired F-actin organization in CNM-neurons

As dendritic spine morphology critically depends on the underlying actin network (Okamoto, et al., 2009; Hotulainen, et al., 2010; Matus, et al., 2000) and actin dynamics is significantly affected in the CNM context (González-Jamett, et al., 2017) we next evaluated actin organization in WT and HTZ neurons. Hippocampal neurons primary cultured from neonatal mice (P0) were fixed, stained with the F-actin-binding toxin phalloidin-rhodamine-B and visualized by confocal microscopy. The phalloidin fluorescence intensity inside dendritic spines was quantified as an estimation of the F-actin content. As shown in Figure 6, the phalloidin-fluorescence intensity was significantly lower in HTZ compared to WT dendritic spines (Figure 6a-b) suggesting defects in the underlying spine-actin network. Indeed, HTZ cultured hippocampal neurons stained with phalloidin exhibited a significantly higher frequency of longer spines and lower head/spine ratio compared to WT neurons (Figure c-e) supporting the idea that actin-dependent dendritic spine features could be impaired in the CNM context. As the relative amounts of filamentous (F-) and monomeric (G-) actin is a determinant factor influencing dendritic spine remodeling (Okamoto, et al., 2009), we quantified the F/G actin ratio in the hippocampus and cortex of WT and HTZ mice. Our results showed that the balance between the F- and G-actin is perturbed in brains of HTZ compared to WT mice (Figure supplement S5) further suggesting that synaptic defects in HTZ mice could rely on disturbances in the actin dynamics.

**Figure 6.**
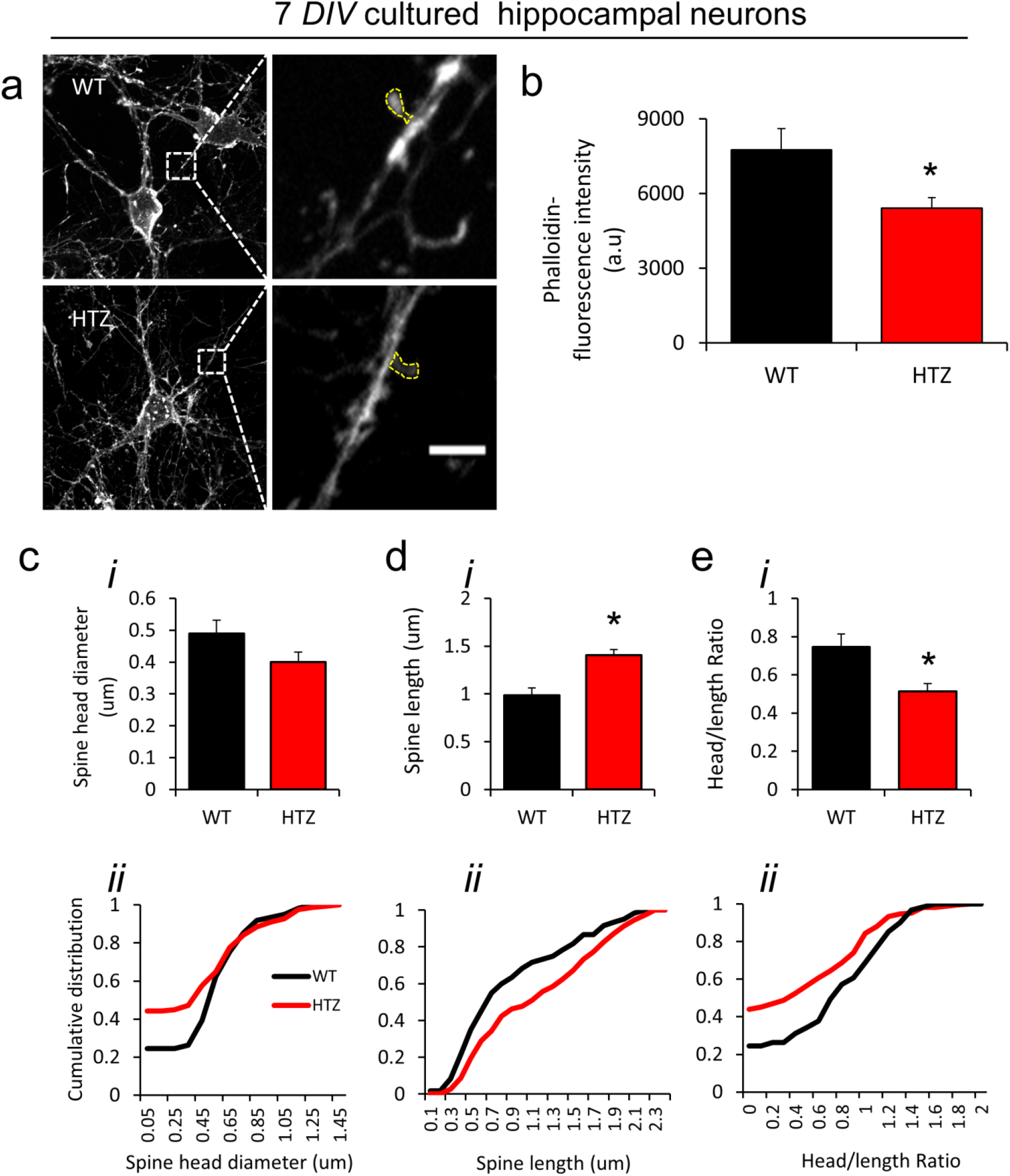
Reduced phalloidin-reactivity in dendritic spines of HTZ neurons. 7 *DIV* hippocampal neurons, primary cultured from WT and HTZ pups (P0), were fixed, stained with the F-actin binding toxin phalloidin-rhodamin-B and visualized by confocal microscopy using comparable acquisition parameters **(a)** Representative phalloidin-stained neurons and dendritic shafts per genotype. Yellow dotted lines delimit dendritic spines inside which phalloidin-fluorescence intensity was measured. Scale bar= 2 µm. **(b)** Mean phalloidin-fluorescence intensity in WT (black bar) and HTZ (red bar) dendritic spines. N=61 WT spines and 166 HTZ from 3 WT and 18 HTZ neurons. Bars are mean ± SEM *p<0.0002 compared to WT condition, Mann-Whitney 2-tail test for non-parametric data. **(c-e)** Average (i) and cumulative distribution (ii) of spine head diameters (c), spine lengths (d) and head/length ratios (e) in WT and HTZ phalloidin-stained neurons. N=61 WT spines, 166 HTZ spines. *p<0.005 versus WT condition, Mann-Whitney 2-tail test for non-parametric data.

All together the results presented here demonstrate that dynamin-2 is an important regulator of the excitatory synaptic transmission in the mammalian brain by modulating actin-dependent dendritic spine structure. This dynamin-2 function appears to be perturbed by the p.R465W mutation, impairing synaptic structure and function and consequently leading to cognitive defects in the CNM context.

## Discussion

Dynamin-2 is a large GTP-ase, member of the dynamin superfamily, that regulates membrane remodeling and actin dynamics, orchestrating actin and membrane-based processes such as exocytosis (González-Jamett, et al., 2013; Shin, et al., 2018), endocytosis (Van der Bliek, et al., 1993; Damke, et al., 1994; Loerke, et al., 2009) and endosomal recycling (Nicoziani, et al., 2000) among others.

In the nervous system, dynamin-2 is expressed at the pre-and post-synaptic levels (Okamoto, et al., 2001). At the pre-synapses dynamin-2 participates in the endocytic recycling of SVs, allowing the resupply of the ready-releasable pool in response to high frequency neuronal activity (Tanifuji, et al., 2013). At the post-synapses dynamin-2 regulates the availability of neurotransmitter receptors at the surface membranes (Carroll, et al., 1999; Kabbani, et al., 2004; Bhatnagar, et al., 2001; Wang, et al., 2017). Of particular relevance for excitatory synaptic transmission is the role played by dynamin-2 in AMPAR trafficking. The GTP-ase activity of dynamin-2 is required for the insertion of AMPAR into spines, by mediating their lateral diffusion from dendritic shafts to PSDs (Jaskolski, et al., 2009). In addition, dynamin-2 GTP-ase activity has been implicated in the removal (Carroll, et al., 1999; Chowdhury, et al., 2006) and recycling (Lu, et al., 2007; Zheng, et al., 2015) of AMPARs from and to PSDs. Hence, as AMPAR mediates the majority of fast excitatory synaptic transmission in the mammalian brain (Anggono, et al., 2012a), dynamin-2 could play a key regulatory function, supporting synaptic transmission and plasticity. In agreement with this idea, pharmacological inhibition of dynamin’s GTP-ase activity with dynole 34-2 (Arriagada-Diaz, et al., 2020) or dynasore (Fa, et al., 2014) significantly reduces LTP in hippocampal slices. Here, we demonstrate that the CNM-causing p.R465W mutation negatively impacts on the synaptic function of dynamin-2, leading to defects in excitatory synaptic transmission and plasticity in the brain of HTZ mice. As dynamin-2 is mostly expressed at the post-synaptic level (Okamoto, et al., 2001) it is likely that these HTZ defects in synaptic transmission rely on post-synaptic mechanisms such as AMPAR trafficking. Indeed, we observed a reduction in the magnitude of LTP in HTZ slices when we induced potentiation both electrically as well as when we did it chemically with glycine. As the chLTP protocol elicits potentiation of the synaptic strength by directly activating NMDARs without mediating the presynaptic release of glutamate (Lu, et al., 2001; Zhang, et al., 2014), these results are consistent with major post-synaptic mechanisms underlying HTZ synaptic defects. In fact, the PPF index was unchanged between WT and HTZ hippocampal slices (Figure 3d). As PPF is an estimation of the neurotransmitter release probability (Zucker, et al., 2002) it is unlikely that the changes that we observed in excitatory synaptic transmission in HTZ hippocampal slices are related to changes in the glutamate release probability. This is agreement with a recent report of Moro and collaborators who found that dynamins are dispensable for synaptic vesicle exocytosis (Moro, et al., 2021).However, we cannot rule out presynaptic defects in the CNM context. The fact that the amplitude of the fiber volley was significantly higher in HTZ compared to WT slices (Figure 2) could mean that more presynaptic terminals are recruited in order to compensate a reduction in the SV releasable pool. As dynamins’s GTP-ase activity of dynamin-2 regulates SV recycling (Tanifuji, et al., 2013; Watanabe, et al., 2013; Cheung, et al., 2019) it is possible to visualize a scenario in which the p.R465W mutation in dynamin-2 affects the size of the SV releasable pool, impacting neurotransmission. More experiments are needed to address this possibility.

Regardless of whether defects were pre or post-synaptic, there was a significant impairment in synaptic transmission and plasticity in HTZ compared to WT brains. Congruently, HTZ mice exhibit a cognitive decline, manifested as defects in recognition and spatial memory which can be explained by perturbations in the synaptic function. In this regard, even though CNM primarily affects skeletal muscles (Romero, et al., 2011; González-Jamett, et al., 2017) learning disabilities and limited intelligent quotient have been reported in dynamin-2-linked CNM patients (Jeannet, et al., 2004; Fischer, et al., 2006; Echaniz-Laguna, et al., 2007; Böhm, et al., 2012). As we observed here in HTZ mice, cognitive defects manifested by CNM patients could be due to disturbances in the synaptic function. Remarkably, in addition to changes in the synaptic strength, we also observed structural modifications in dendritic spines of the HTZ compared to WT neurons. In this regard, dendritic spine density was significantly lower in pyramidal neurons from adult HTZ mice (Figures 4-5). As the majority of the synaptic contacts between excitatory neurons are made on dendritic spines (Harris, et al., 1994) a net reduction in spine density can lead to significant defects in synaptic transmission, such as those observed in HTZ hippocampal slices. Moreover, dendritic spine morphology, which is closely related to the maturity and functionality of synapses (Holtmaat, et al., 2009; Gipson, et al., 2017), also underwent modifications in HTZ compared to WT neurons (Figures 4, 5 and 7). Although on average we did not observe significant differences in spine morphology between genotypes, the cumulative distribution of the spine length, spine head diameter and head/length ratio yielded important variations. Whilst mature HTZ hippocampal neurons exhibited a higher frequency of shorter spines, neonatal hippocampal neurons cultured from HTZ pups showed a higher frequency of immature larger spines with a significant reduction in the head diameter/spine length ratio. Moreover, cortical neurons from adult HTZ brains exhibited higher frequency of shorter spines, with smaller heads compared to WT neurons. As spines with large head and large head/spine ratio have been found to correlate with an increased PSD area and AMPAR density (Matsuzaki, et al., 2001) a reduction in spine head diameter or the head/length ratio could explain defects in synaptic transmission and plasticity.

**Figure 7:**
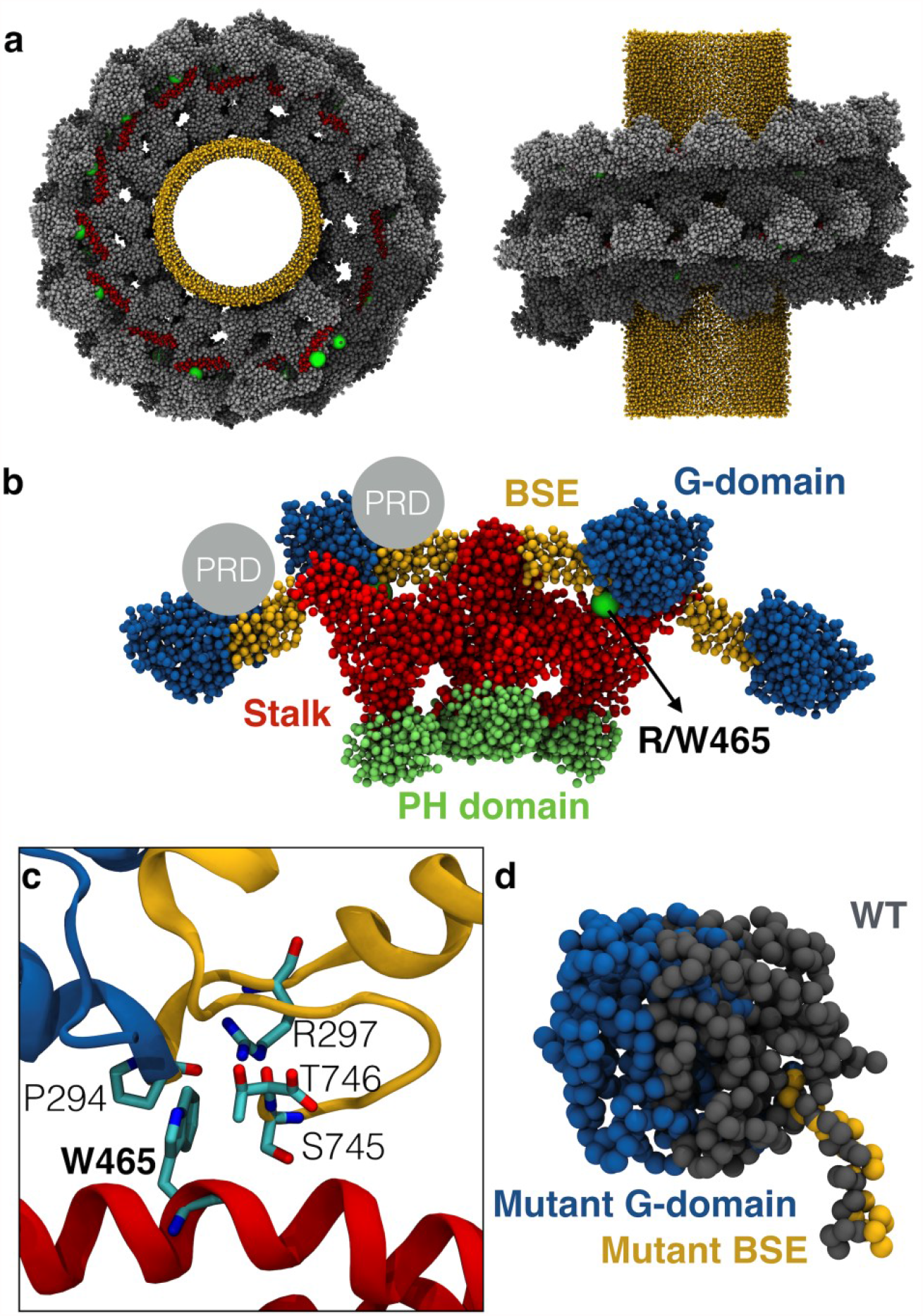
The CNM-causing p.R465W mutation localizes near to a putative direct actin-binding motif in dynamin-2 **(a)** Corse-grained model of a dynamin-2 helix surrounding a lipid nanotube. Dynamin-2 is represented in silver and gray; Putative actin-binding region proposed by Gu et al., (2010) (residues 399 to 444 in the middle domain) are shown in red; the residue R/W465 is represented in green and the lipid nanotube in yellow. **(b)** Tetramer configuration in the dynamin-2 helix. G-domains are colored in blue, BSEs are represented in yellow, stalks are shown in red and PH domains in lime. The position of PRD is represented only in two dynamins-2 as a silver circles. Green dot represent R/W465. **(c)** Amino acids of the BSE that are close to the R/W465. This interaction occurs in a zone close to the PRD. **(d)** G-domains and BSE of WT and mutant dynamin-2. The extension of BSE in the mutant dynamin-2 changes the position of the G-domain.

A major factor regulating spine morphology is the actin cytoskeleton (Okamoto, et al., 2004; Bosch, et al., 2014). A dynamic F-actin network exists in dendritic spines, which acts primarily as a cytoskeleton component defining spine architecture, but also as a scaffold for the recruitment of post-synaptic proteins including AMPAR (Okamoto, et al., 2009; Cingolani, et al., 2008; Stefen, et al., 2016).Therefore, remodeling of the spine-actin is critical for structural and functional synaptic plasticity (Kim, et al., 1999; Okamoto, et al., 2004) and it is not surprising that perturbations in the actin dynamics lead to different synaptopathies (Penzes, et al., 2012; Pelucchi, et al., 2020). In fact, the changes that we observed in the density and shape of dendritic spines in HTZ neurons (Figures 4-6) are compatible with defects in actin dynamics. In this regard, we previously demonstrated that the actin organization and remodeling are disrupted in muscle cells of HTZ mice (González-Jamett, et al., 2017) suggesting that a similar mechanism could operate in central synapses. Our results here suggest that an imbalance in the relative amounts of F- and G-actin occurs in the brain of HTZ mice (Figure supplement S5) which could be associated with a defective spine-actin dynamics. An increase in the F/G ratio as we observed in HTZ compared to WT brains (Figure supplement S5) could be related to a less dynamic neuronal actin network, with a reduced ability to remodel itself to support synaptic transmission. But, how the p.R465W mutation could impact on dynamin-2 function leading to defects in actin and actin-based mechanisms? The residue 465 localizes near to a putative direct actin-binding motif in dynamin-2 (Gu, et al., 2010, red region in Figure 7a). As these authors proposed, unassembled dynamins could bind to short actin filaments via their middle domain and in turn oligomerize in rings promoting actin polymerization (Gu, et al., 2010; Gu, et al., 2014; Shiffer, et al., 2015). If that is true, the substitution of a positive-charged arginine by a non-polar tryptophan in the oligomerization-middle domain of dynamin-2 could certainly influence its actin-binding properties affecting actin dynamics. In this regard, by using full-atom and coarse-grained molecular simulations, we previously demonstrated that the p.R465W mutation affects the interaction between the residue 465 and surrounding amino acids, modifying the conformation of the BSE and consequently impacting on the GTPase-domain position and dynamin-2 self-assembly (Hinostroza, et al., 2020). Recent evidences suggest that helix of dynamin-2 interact with F-actin via their outer-rim (Zhang, et al., 2020). This region includes the C-terminus PRD that is adjacent to the GTPase domain and to the BSE (Figure 7b). It is noteworthy that the R/W 465 residue interacts with the BSE in a region that is close to the starting point of the PRD (Figures 7b-c). As the p.R465W mutation in dynamin-2 extends the angle between the G domain and the BSE (Hinostroza, et al., 2020) in the tetramer and helical configuration (Figure 7d) we speculate that the arginine/tryptophan substitution in the residue 465 not only modifies the BSE conformation but also changes the position of the PRD, being able to affect the PRD-actin interaction. This assumption is in agreement with a model in which the actin-remodeling activity of dynamin-2 is disrupted by CNM-causing mutations. In accordance with this idea, Lin and collaborators recently demonstrated that dynamin-2 regulates the bundling of actin at the postsynaptic membranes of neuromuscular junctions (NMJs). Remarkably, CNM-linked mutations seem to disrupt this dynamin-2 function impairing actin remodeling and perturbing postsynaptic structures at the NMJs (Lin, et al., 2020). A similar mechanism could be operating at central excitatory synapses in the brain of HTZ mice, explaining the defects that we observed in dendritic spine morphology, synaptic transmission and cognitive functions.

In our knowledge, this is the first report addressing the mechanisms that lead to cognitive defects in dynamin-2-linked CNM. Our data support a model in which the actin cytoskeleton organization is impaired, leading to structural and functional synaptic defects. In addition, our results contribute to the better understanding of the still scarce knowledge about the function of dynamin-2 at central synapses.

## Methods

### Animals

HTZ mice harboring the p.R465W mutation in dynamin-2 is a mammalian model of CNM. These mice recapitulate most of the CNM-signs, which start at 1 mo and progress as animals age (Durieux et al., 2010). HTZ and WT littermates (C57BL/6 strain) were housed at room temperature with *ad libitum* access to food and water. Mice were maintained on light-darkness cycles of 12–12 h, according to standard protocols. Genotyping was performed by PCR as previously described (Durieux et al., 2010) using DNA extracted from ears. Primers used were: 3′-CTGCGAGAGGAGACCGAGC-5′ (forward) and 3′-GCTGAGCACTGGAGAGTGTATGG-5 (reverse). PCR products were electrophorated in agarose gels and bands of 445 bp and 533 bp, representing the WT and p.R465W mutated allele respectively, were detected. All animal protocols described here were conducted in accordance with the approved protocols of the Institutional committee for the care of laboratory animals of Universidad de Valparaíso (BEA131-18).

### Behavioral tests

Recognition and spatial learning and memory were assessed in 6 m.o WT and HTZ mice by application of the Novel Object Recognition (NOR) test and the Barnes maze test (BM), respectively. Locomotion capabilities in mice were evaluated by application of a standard Open Field (OF) test. All behavioral tests were carried out in a room conditioned for this purpose, with controlled temperature (21 ° ± 2 ° C), white noise (50 dB) and regulated luminosity (200 lux).

#### NOR

This test is based on the spontaneous tendency of rodents to spend more time exploring a novel object than a familial one. Maze consisted of a 50 cm x 40 cm x 63 cm square white acrylic box in which mice explored freely during 5 min throughout three phases: (i) sample, where mice explored a pair of identical objects; (ii) retention, where mice were removed for cleaning and changing objects; and (iii) choice, where mice explored a pair of different objects: a familiar object (F) and a new object (N). Each session was repeated during 3 days. The time that mice spent exploring F or N was quantified. A recognition index (D1) was calculated as [N exploration time-F exploration time.

#### BM

This task is dependent on the intrinsic inclination of rodents to escape from an aversive environment (Barnes 1979). It consists in a 70 cm diameter white circular platform elevated 90 cm from the floor with 20 equally spaced holes along the perimeter (each 7 cm in diameter) located 2 cm from the edge of the platform. The animals are oriented by means of spatial clues located on the walls of the room, in order to find and remember the position of a black plexiglass escape box (17 × 13 × 7 cm) that is hidden under one of the holes. The maze was illuminated with 4 incandescent lights to produce a light level of around 600 lux fall on the circular platform. BM test consisted of 9 session days, 4 trials per session along which three phases were carried out: (i) pre-training phase, where mice were trained to find the escape platform (ii) training phase, where mice remember the position of the escape box guided by the spatial clues arranged in the room, (iii) reversion phase, where the location of the escape box was changed but maintaining the position of the spatial clues to evaluate memory flexibility. In all phases, if mice did not find the target hole and did not enter into the escape box during 3 minutes, the test was terminated. Subsequently, the experimenter guided the animal to the escape box and allowed it to remain in the box for 1 minute before returning to their holding box. Latency or time that took to animals to find the target hole and enter into the scape box was quantified.

#### OF

Mice were placed in the center of a 50 cm x 40 cm x 63 cm white acrylic box and left to explore freely during 5 minutes, along 3 sessions (1 session per day). Total distance traveled and the percentage of walking time was quantified.

### Field electrophysiological recordings

To evaluate basal strength of synaptic transmission and synaptic plasticity we evoked field excitatory postsynaptic potentials (fEPSPs) in WT and HTZ hippocampal slices by stimulating the Schaffer Collaterals (CA3) and recording the synaptic responses in *stratum radiatum* of CA1. Hippocampal slices were prepared as previously described (Ardiles et al. 2014). Animals were euthanized under deep anesthesia using isoflurane at saturation and their brains were quickly removed after craniotomy. Brains were sectioned in two hemispheres, removing hippocampus to be sectioned in 300 µm thick slices and immersed in dissection buffer (in mM: 212.7, sucrose 26 NaHCO_3_, 1.23 NaH_2_PO_4_, 10 D-glucose, 5 KCl, 2 CaCl_2_, 1 MgCl, 3 pyruvate) using vibratome (Vibratome 1000 plus, Ted Pella Inc., CA, USA). Slices were transferred and kept for 1 hour at room temperature in artificial cerebrospinal fluid (ACSF, in mM: 124 NaCl, 26 NaHCO_3_, 1.23 NaH_2_PO_4_, 10 D-glucose, 5 KCl, 2 CaCl_2_, 1 MgCl, 3 pyruvate). All recordings were obtained in an immersion chamber perfused with ACSF (30 ± 0.5°C; 2 ml / min). Stimulation was performed with pulses of 0.2 ms duration, administered through concentric bipolar stimulation electrodes. Basal responses were recorded using mean stimulation intensity compared to maximum. Basal synaptic transmission was analyzed by determining input-output relationship (I /O) of the fEPSP generated by gradually increasing the stimulation intensity. Input corresponded to the amplitude of the fiber volley and the output to the initial slope of the fEPSP. To evaluate long-term potentiation (LTP), slices were subjected to a high-frequency electrical stimulation protocol or a chemical stimulation protocol. LTP using high-frequency electrical stimulation was performed using a standard theta burst (TBS) protocol, which consists of 4 trains at 100 Hz for 1 min (Flores-Muñoz et al. 2020; Gajardo et al. 2018). On the other hand, chemical LTP (chLTP) was induced by perfusing a glycine solution (glycine, Gly 600 µM + *picrotoxin*, PTX 50 µM + strychnine Stric 3 µM) (X. Y. Zhang et al. 2014), directly on the hippocampal slices during 5 minutes. Gly is co-agonist of NMDAR, PTX is an inhibitor of the inhibitory

GABAR-dependent transmission and Stric is an inhibitor of Gly-receptors. Once the application time of solution had elapsed, it was removed and the registration buffer was replaced with a new one. In both cases, fEPSPs were recorded for 20 minutes prior to stimulation and for 50 minutes after the application of the electrical stimulus (TBS) or chLTP.

To assess long-term depression (LTD), a standard low-frequency electrical protocol was induced which consists of a 1 Hz stimulation for 15 minutes (Gajardo et al. 2018). fEPSPs were recorded for 20 minutes prior to stimulation and for 60 minutes after the application of LFS.

Paired pulse facilitation (PPF) was obtained by stimulating at different intervals in a range between 25-300 ms and recording excitatory post-synaptic potential (EPSP). A PPF index was estimated by dividing the amplitude of the second response over the first one (R2/R1).

### Spine density and morphology

HTZ mice over 6 mo and age-matched WT littermates were sacrificed, brains were quickly removed and processed using the FD Rapid Golgi-Stain-TM kit (FD Neuro Technologies) according to the manufacturer instructions. Golgi impregnated cortical and hippocampal neurons were imaged using a Leica DM500 microscope equipped with a camera system, a 63X /1.40 oil (HCPL Apo, Leica) objective and the Leica acquisition software. Pyramidal neurons were analyzed and processed using the ImageJ software (NIH, USA). To evaluate the spine density and morphology, measurements were performed in dendritic segments of 10 to 20 μm, visualized from four apical dendrites per neuron. Dendritic-protrusions were categorized following the parameters revised by Risher and collaborators (Risher et al., 2014). Long, thin without-head protrusions of more than 2 um long were classified as immature filopodia; wide-head (>0.6 um width) and short protrusions (<1 um) were classified as mature mushroom-spines; wide-head/without neck protrusions (length: width ratio <1 um) were classified as stubby-spines; thin and short (<2 um) headed protrusions were classified as thin-spines. Cup-shaped protrussions were classified as branched spines (Figure supplement S6). Measurements were made by two different experimenters, blind to the genotype of the samples.

### Neuron primary culture

Hippocampal neurons were primary cultured from post-natal mice (P0) following the protocol described by Beduoain and collaborators (Beduoain et al., 2012). Briefly, hippocampi from individual pups were dissected, trypsinized for neurons dissociation and plated in poly-L-lysine treated coverslips and maintained in medium containing B27-supplement (GIBCO), glutamax (GIBCO) and Penicilin/Streptomycin mixture of antibiotics (GIBCO). Genotype of each single culture was confirmed a *posteriori* from tail-segments of pups. 2 days after culture neurons were treated with 5µM cytosisne-arabinoside (Ara-C) to inhibit the proliferation of non-neuronal cells. At 7 *DIV*, neurons were fixed with 4%PFA + 4% sucrose during 10 min at 37°C, permeabilized with 0.1% triton-X100 in saline phosphate buffer (PBS, mM: 137 NaCl, 2.7 KCl, 10 Na_2_HPO_4_, 2 KH_2_PO_4_, pH 7.4) during 10 min at room temperature and stained during 1 h with 1µM phalloidin-Rhodamin-B (Sigma) before confocal visualization. Images were acquired in a confocal microscope (upright Eclipse Nikon 80i) using a 60x magnification objective. Single confocal images were acquired with the EZ-C1 (Nikon) software using similar acquisition settings between comparable samples. Analyses of dendritic spines stained with phalloidin were made with the ImageJ software (NIH, USA) by two different experimenters, blind to the genotype of the samples. The phalloidin fluorescence intensity was measured inside spines, without including the neighboring dendritic shaft.

### F/G actin assay

Hippocampal and cortical tissue from WT and HTZ mice were lysed and homogenized in conditions that stabilize filamentous (F-) and monomeric (G-) actin using an F/G actin commercial assay (Cytoskeleton Inc.). Lysed extracts were ultracentrifuged at 100,000 g for 1 h at 37 °C in order to separate pellets (F-actin) and soluble (G-actin) fractions. The F-actin pellet was resuspended in a depolymerizing buffer (Cytoskeleton Inc) and then F-and G-actin fractions were diluted in loading buffer (50 mM Tris–HCl, 2% SDS, 10% glycerol, 1% beta-mercaptoethanol and bromophenol blue). Samples were subjected to electrophoresis in 12%-SDS-PAGE, transferred to a PVDF membrane, blocked for 1 h with 5% non-fat milk in TBS-T (mM:150 NaCl, 50 Tris/HCl, pH 7.4, 0.05% Tween-20), incubated with a polyclonal antibody against actin (1:500; Cytoskeleton Inc.) and developed by chemiluminescence. Densitometric analysis of the F- and G-actin bands was performed with the ImageJ software.

### Statistical analyses

GraphPad Prism 9 software was used to perform the statistical analyses. Paired or unpaired non-parametric Mann-Whitney test were used to compare WT and HTZ conditions. Two-way ANOVA followed by Bonferroni multiple comparison post-test was also used in electrophysiological experiments. Data in graphs represent the mean ± SEM, with a p-value <0.05 being significant. If not specified, differences were not statistically significant.

## Acknowledgements

We thank Dr. Isaac García-Carrillo (Escuela de Odontología, Universidad de Valparaíso) for facilitating his laboratory for western blot development. This work was supported by the grants Fondecyt 11180731 (to AMG-J), Fondecyt 1201342 (to AOA), and ICM-ANID ICN09-022 CINV (to AMG-J, AOA, and AMCD) and by ANID scholarship 22200154 (to JA-D).

---

**Figure S1:**
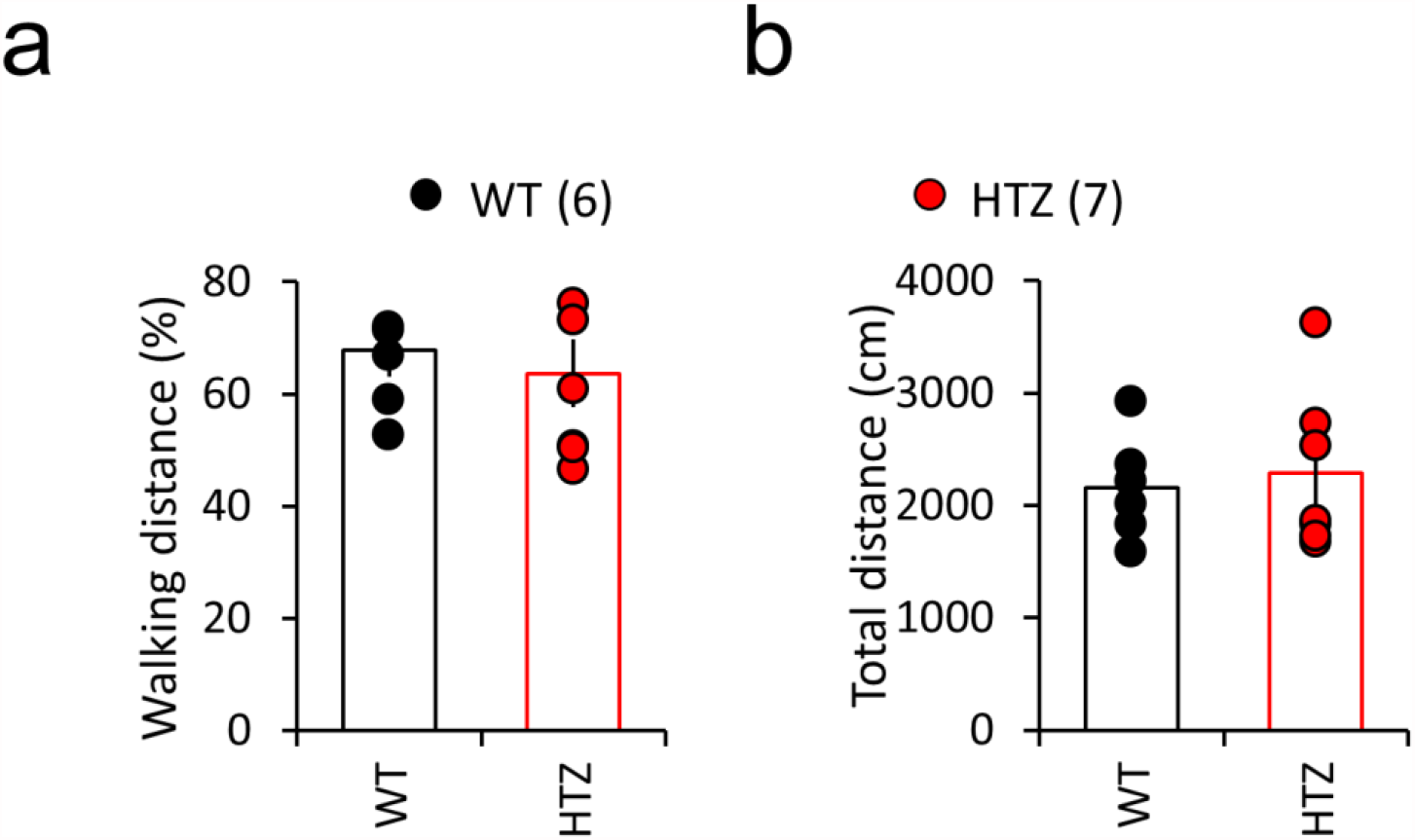
Locomotion activity is not different between WT and HTZ mice. WT (black circles) and HTZ (red circles) mice over 6 m.o were left to explore freely in an open field arena during 5 min. (a) Mean percentage of the walking time and (b) total distance traveled are plotted per each experimental group. All data are represented as mean ± SEM. N= 6 WT and 7 HTZ mice. Statistical differences were calculated using a non-parametric Mann-Whitney test.

**Figure S2:**
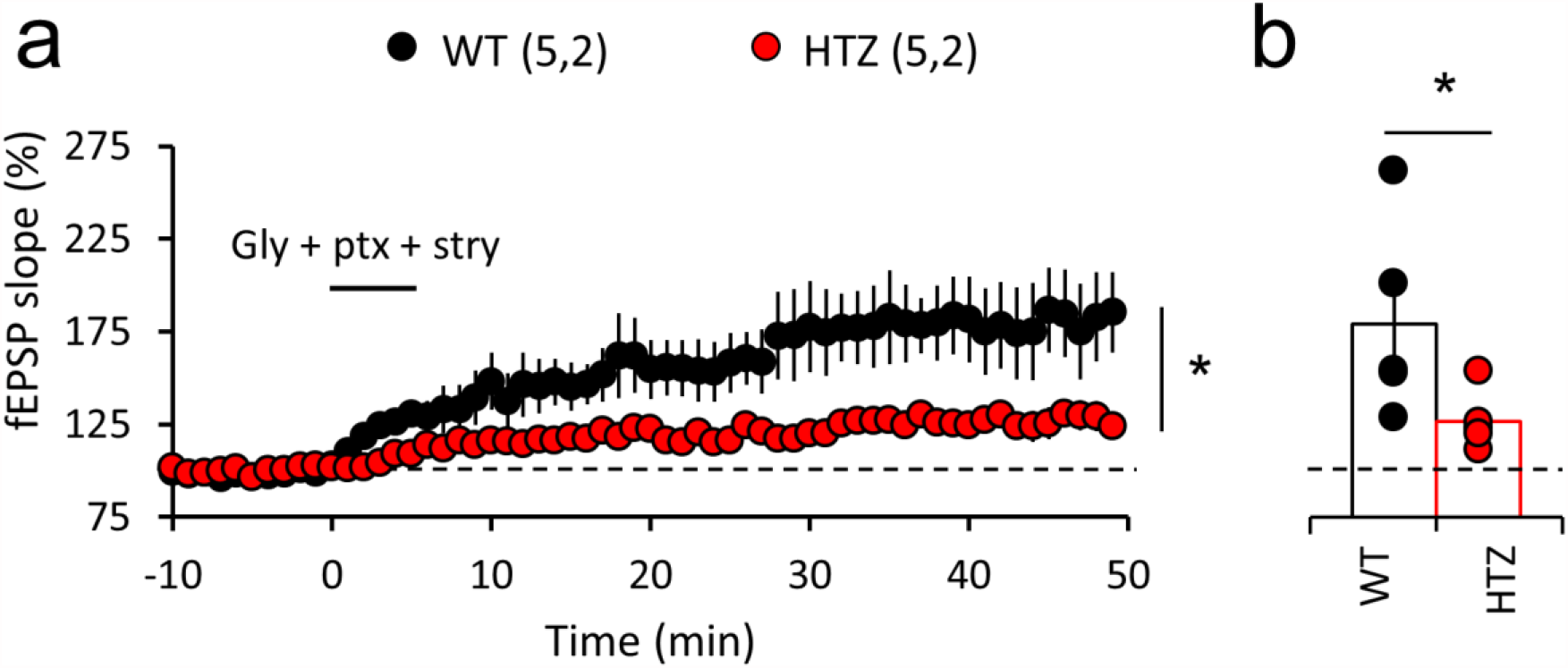
Impaired glycine-induced LTP in HTZ hippocampal slices. **(a-b)** Hippocampal slices from 6 m.o WT (black circles) and HTZ (red circles) mice were stimulated with the application of 600 µM glycine + 50 µM PTX + 3 µM strychnine in order to chemically induce LTP and fEPSPs were recorded during 50 min. 2-Way ANOVA, F_(1,480)_ = 208.2, *p <0.0001 followed by Bonferroni post hoc test (p < 0.05) compared to the WT condition. Note that HTZ hippocampal slices exhibit a significantly reduced potentiation at the end of the recording (b) compared to the WT slices (N= 5 slices, 2 WT mice; 5 slices, 2 HTZ mice). Mann-Whitney test * p <0.005 compared to the WT condition. All data are represented as mean ± SEM.

**Figure S3.**
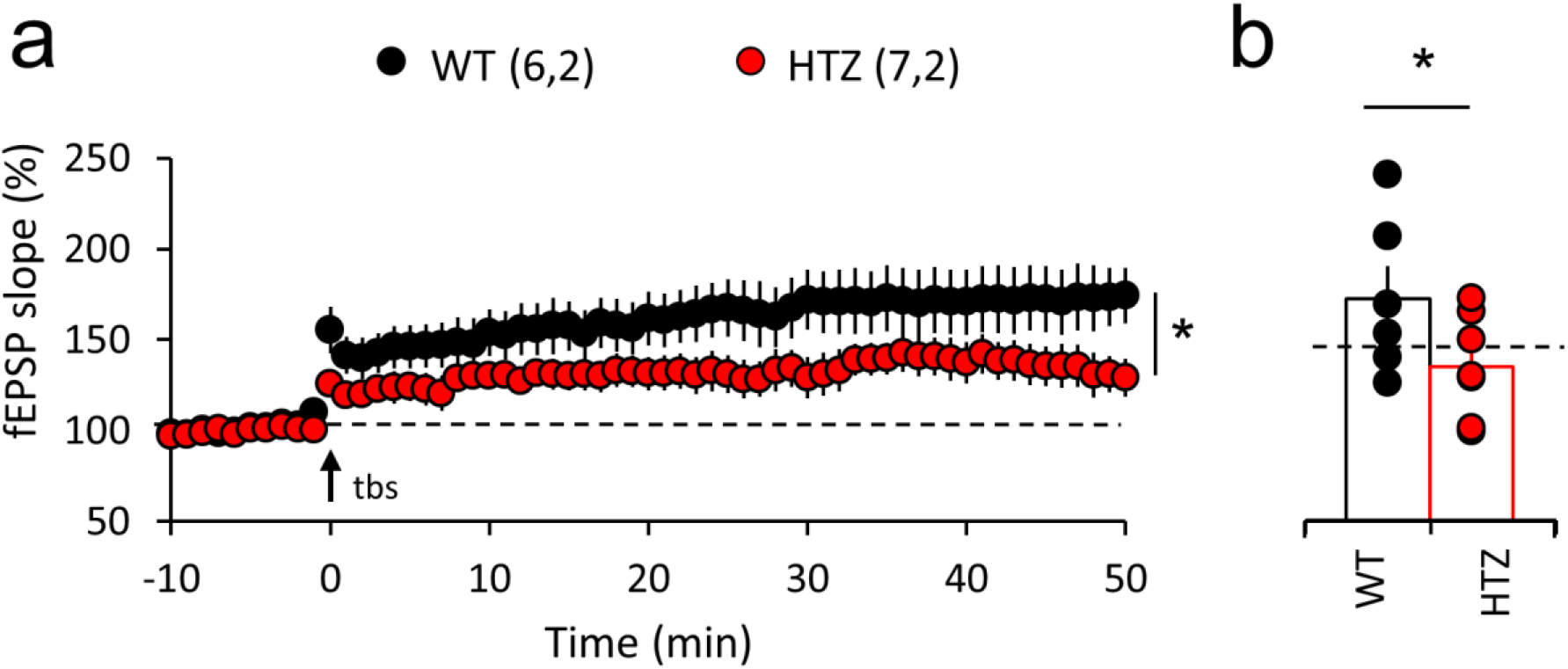
LTP impairment in HTZ hippocampal slices is independent of inhibitory synaptic transmission. **(a-b)** Hippocampal slices from 6 m.o WT (black circles) and HTZ (red circles) mice were stimulated with a standard TBS protocol in the presence of the GABAR-inhibitor Picrotoxin (PTX). fEPSPs were recorded during 50 min after TBS-application. 2-Way ANOVA, F_(1,671)_ = 245.8, *p <0.0001 followed by Bonferroni post hoc test (p < 0.05) compared to the WT condition. Note that even in the presence of PTX, HTZ hippocampal slices exhibit a significantly reduced potentiation compared to WT slices, suggesting that inhibitory transmission is not involved in the synaptic plasticity defects in HTZ slices. N= 6 slices, 2 WT mice/ 7 slices, 2 HTZ mice). Mann-Whitney test * p <0.005 compared to the WT condition. All data are represented as mean ± SEM.

**Figure S4.**
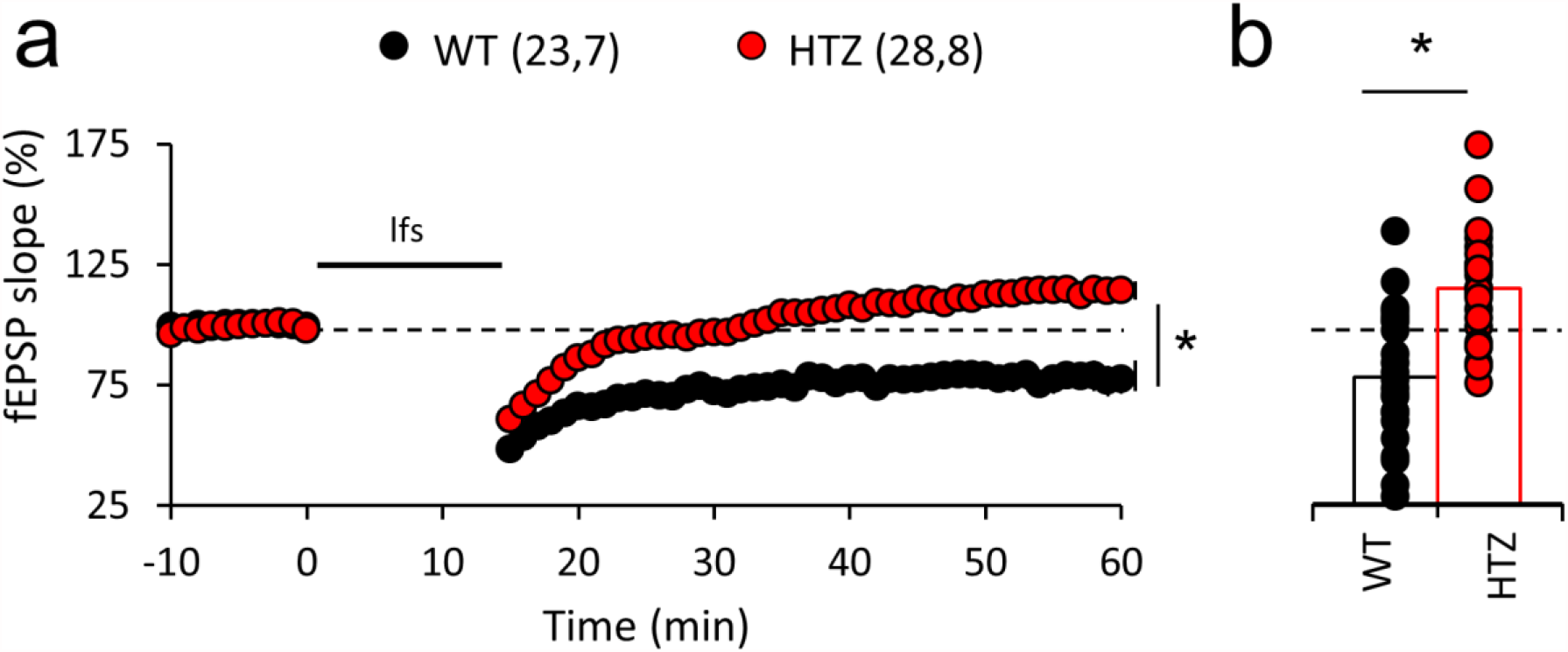
Impaired LFS-induced LTD in HTZ hippocampal slices. **(a-b)** Hippocampal slices from 6 m.o WT (black circles) and HTZ (red circles) mice were stimulated with a standard low-frequency electrical stimulation (LFS) protocol and fEPSPs were recorded during 60 min to estimate depression of the synaptic response (LTD). 2-Way ANOVA, F_(1,2793)_ = 1039, *p <0.0001 followed by Bonferroni post hoc test (p < 0.05) compared to the WT condition. Note that HTZ hippocampal slices exhibit a significantly reduced capability to depress the synaptic response at the end of the recording compared to WT slices (WT: n = 23 slices, 7 mice; HTZ: n = 28 slices, 8 mice). Mann-Whitney test * p <0.005 compared to the WT condition. All data are represented as mean ± SEM.

**Figure S5:**
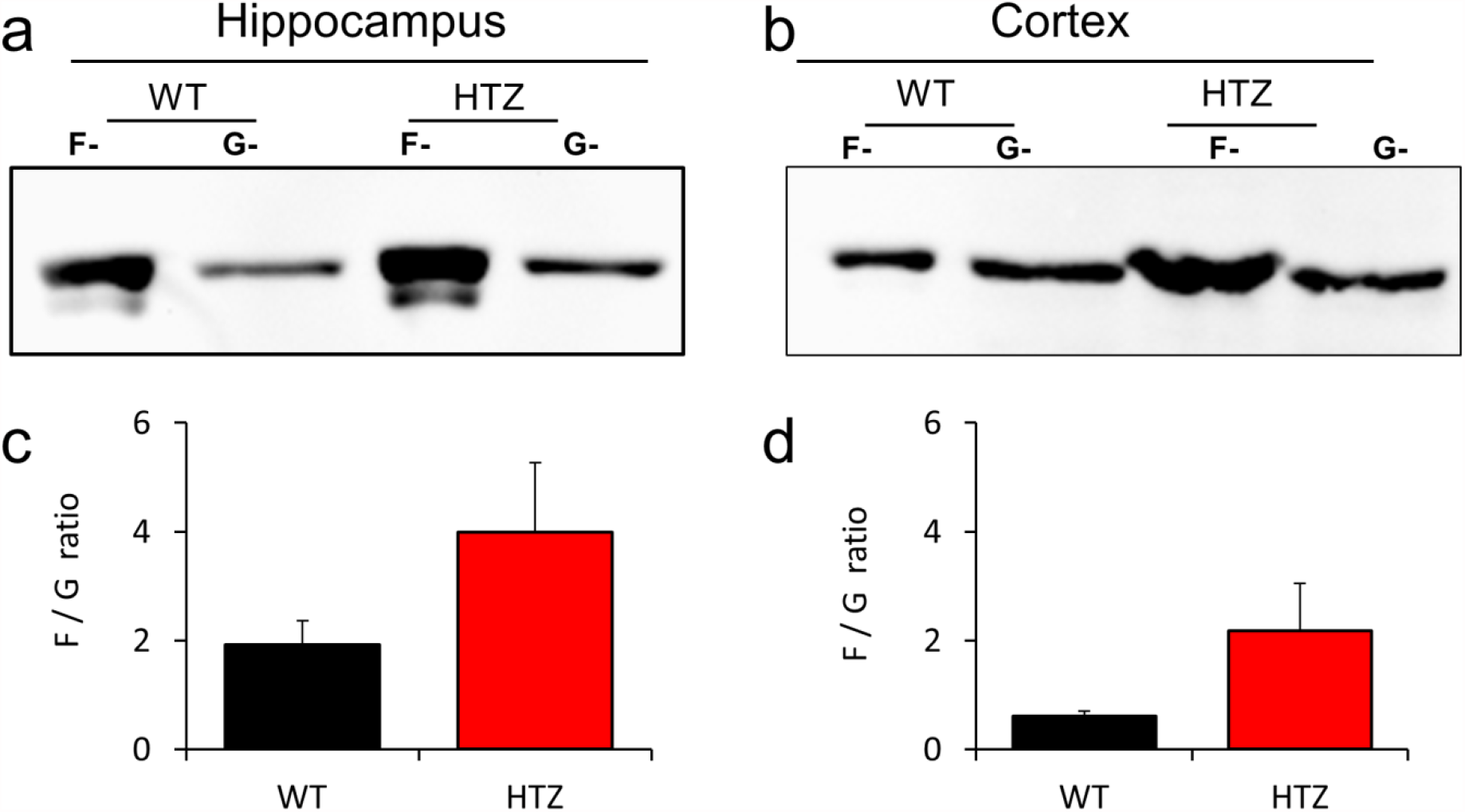
Imbalance in the F/G actin ratio in brains of HTZ mice. Representative blots of the relative amounts of F- and G-actin in the hippocampus (a) and cortex (b) of WT and HTZ mice are shown. (c-d) Graphs summarize the quantification of the densitometry analysis of the F/G actin ratio in hippocampus and cortex (d) isolated from WT and HTZ mouse brains. Bars are mean ± SEM of tissues from 3 WT and 3 HTZ mice. Statistical comparison was done using Mann-Whitney test for non-parametric data. p=0.200 and 0.400 for hippocampus and cortex, respectively. Although not significant, HTZ brain tissues tended to exhibit an increased F/G ratio compared to the WT condition.

**Figure S6.**
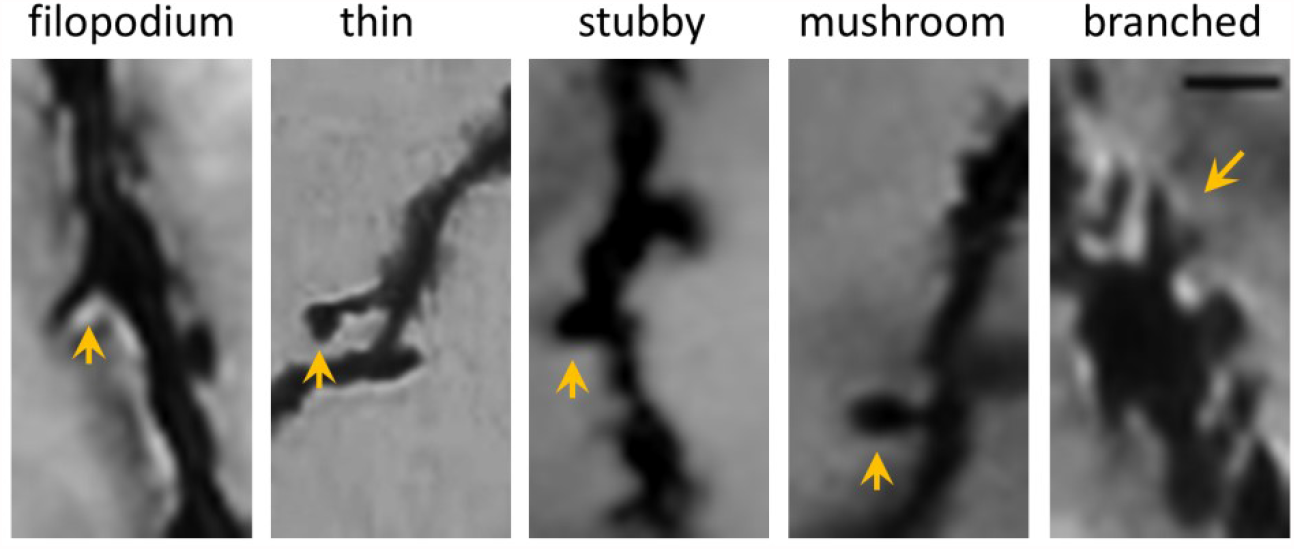
Representative dendritic spine-morphologies. Filopodia are long protrusions (> 2 um) without head. Thin-spines are short (<2 um) protrusions with small heads (<0.5 um). Stubby-spines are wide-head protrusions without neck. Mushroom spines are short protrusions (<1 um) with wide-heads (> 0.6 um width). Branched spines are short cup-shaped protrussions. Yellow arrows point dendritic spines of each morphology. Scale bar=2 um

